# PROTEOME-WIDE ANALYSIS OF CD8+ T CELL RESPONSES TO EBV REVEALS DIFFERENCES BETWEEN PRIMARY AND PERSISTENT INFECTION

**DOI:** 10.1101/327312

**Authors:** Calum Forrest, Andrew D. Hislop, Alan B Rickinson, Jianmin Zuo

## Abstract

Human herpesviruses are antigenically rich agents that induce strong CD8+T cell responses in primary infection yet persist for life, continually challenging T cell memory through recurrent lytic replication and potentially influencing the spectrum of antigen-specific responses. Here we describe the first lytic proteome-wide analysis of CD8+ T cell responses to the Epstein-Barr gamma1-herpesvirus (EBV), and the first such proteome-wide analysis of primary versus memory CD8 responses to any human herpesvirus. Primary effector preparations were generated directly from activated CD8+ T cells in the blood of infectious mononucleosis (IM) patients by in vitro mitogenic expansion. For memory preparations, EBV-specific cells in the blood of long-term virus carriers were first re-stimulated in vitro by autologous dendritic cells loaded with a lysate of lytically-infected cells, then expanded as for IM cells. Preparations from 7 donors of each type were screened against each of 70 EBV lytic cycle proteins in combination with the donor’s individual HLA class I alleles. Multiple reactivities against immediate early (IE), early (E) and late (L) lytic cycle proteins, including many hitherto unrecognised targets, were detected in both contexts. Interestingly however, the two donor cohorts showed a different balance between IE, E and L reactivities. Primary responses targeted IE and a small group of E proteins preferentially, seemingly in line with their better presentation on the infected cell surface before later-expressed viral evasins take full hold. By contrast, target choice equilibrates in virus carriage with responses to key IE and E antigens still present but with responses to a select subset of L proteins now often prominent. We infer that, for EBV at least, long-term virus carriage with its low level virus replication and lytic antigen release is associated with a re-shaping of the virus-specific response.

## Author Summary (156 words)

Epstein-Barr virus (EBV) is a herpesvirus and an important human pathogen that normally persists for life and induces strong CD8+T cell responses. Primary EBV infection can cause mononucleosis (IM) disease and EBV is associated with malignant lymphocytes cancers and epithelial cells. Here for the first time, we described proteome-wide analysis of CD8+ T cell responses to EBV lytic antigens, and compare the CD8 T cell responses between primary and memory CD8 responses to EBV. The EBV primary CD8 T cells responses targeted IE and a small group of E proteins preferentially. By contrast, the memory CD8 T cells responses from EBV persistent infection are dominated by L proteins. This study can help to understand how human CD8 T cells respond evolves with the long-term virus infection. For EBV at least, we infer that long-term EBV virus carriage with its low level virus replication and lytic antigen release is associated with a re-shaping of the virus-specific response.

## INTRODUCTION

Human herpesviruses are ancient pathogens that have co-evolved with our species and its antecedents over millions of years, producing a finely balanced virus-host relationship in which both primary infection and subsequent life-long virus carriage are typically asymptomatic. By contrast, herpesvirus infections can be life-threatening in individuals with congenital or iatrogenic impairment of the T cell system, illustrating the central importance of cell-mediated immune responses in achieving an equable virus-host balance [1]. Our limited understanding of those responses reflects the antigenic complexity of these large DNA viruses with, depending on the agent, 60 to >150 proteins being expressed during the virus replicative (lytic) cycle. While all of these proteins are potentially immunogenic, they include viral evasins which partially shield infected cells from T cell (especially cytotoxic CD8+ T cell) recognition and whose effects upon the immunogenicity of lytic cycle proteins remain unclear[2]. Determining the range of responses that are induced and identifying those that best mediate host control are seen as key to the development both of prophylactic vaccines against primary infection and of immune therapies targeting chronic infection.

Concerted efforts towards a virus proteome-wide mapping of T cell targets have so far been made for herpes simplex (HSV, an alpha-herpesvirus) using a viral gene expression library [3] and human cytomegalovirus (HCMV, a beta-herpesvirus) using a synthetic peptide library [4]. In each case mapping revealed a range of CD8+ T cell targets straddling the immediate early (IE), early (E) and late (L) temporal phases of the virus lytic cycle, with little evidence that early-expressed evasins had limited late protein immunogenicity. Note, however, that the above studies were focussed on responses seen in long-term virus carriage rather than during initial infection, reflecting the difficulty of identifying subclinical primary infections with these agents. As a result, the relationship between target antigen choice in primary versus memory responses, and by inference the possible influence of virus persistence on target choice, remains to be determined. Here we describe studies on Epstein-Barr virus (EBV), an oncogenic gamma1-herpesvirus, whose spectrum of lytic antigen-induced CD8+ T cell responses has never been rigorously analysed. Importantly with this virus, proteome-wide screening can be conducted both on long-term virus carriers and on individuals in whom primary infection is clinically manifest as infectious mononucleosis (IM).

Previous studies, largely based on IM patients, have shown that EBV is orally transmitted and replicates through lytic infections in the oropharynx, probably involving mucosal epithelium and some locally-infected B cells, leading to high levels of infectious virus being released in the throat [5]. At the same time, the virus spreads within the generalised B cell system through a growth-transforming infection, transiently expressing a spectrum of 8-10 latent cycle antigens. Thereafter, antigen expression appears to be suppressed and the virus is then carried for life as a truly latent (i.e. antigen-null) infection in the recirculating B cell pool[6]. Occasionally individual cells within this pool will reactivate into lytic cycle, and some of those reactivations are thought to re-seed foci of oropharyngeal replication, thereby giving rise to the low levels of infectious virus intermittently detectable in the throat washings of virus carriers[7].

How such lytic and latent infections are perceived by the host T cell system has long been of interest to immunologists, prompted initially by the observation that IM blood contains large expansions of activated CD8+ T cells. Much of this acute primary response appears to be EBV-specific and predominantly focused on lytic antigen targets [8, 9]. Thus we have described examples of particular IM patients where a significant fraction (in extreme cases up to 50%) of the activated CD8+ T cell population is attributable to responses against 1-3 individual EBV peptide epitopes, with smaller responses to other epitopes also detectable [10, 11]. The immunodominant epitopes are usually but not exclusively derived from lytic rather than latent cycle proteins, and in the best studied examples on restriction elements such as HLA-A*0201 and B*0801 those responses mapped to IE or certain E proteins with little evidence of L-specific reactivities [9, 11, 12]. Having defined the relevant peptide epitopes, HLA-peptide tetramers allow one to follow the evolution of specific responses into memory over the ensuing weeks/month. Such studies show that, as the acute disease resolves, the responding cells lose activation markers and are culled in number but, at least in the short term, retain their hierarchy of representation within the circulating CD8+ T cell pool [13]. Moreover, when healthy virus carriers with the same HLA alleles were studied by tetramer staining or epitope peptide stimulation ex vivo, the same reactivities were observed often with a similar hierarchy of inter-epitope representation[14] There were occasional examples where particular reactivities disappeared over time post-IM[13], but overall the results implied that the range of memory responses to EBV lytic cycle antigens and their hierarchy of immunodominance was essentially determined during primary infection.

However the above studies were limited in their scope, mapping responses through a narrow range of HLA alleles against just a subset of lytic cycle antigens, namely the two IE proteins, 11 of 32 E proteins and 10 of 36 L proteins[12]. Whether the apparent IE > E > L hierarchy of immunodominance among lytic cycle proteins as CD8 targets would withstand more complete analysis remained an open question. Interestingly two subsequent studies of CD8+ T cell memory, in macaques carrying the analogous rhesus gamma1-herpesvirus[15] and in healthy EBV carriers[15, 16], focussed on selected L antigen targets and detected occasional strong responses, though whether such responses were equally represented at the time of primary infection was unresolved. The present work, using a proteome-wide approach to study lytic antigen-induced CD8+ T cell responses, provides the first comprehensive survey of EBV antigen choice both in acute IM patients and in long-term carriers, and describes significant differences in that choice between the two phases of infection.

## RESULTS

### EBV lytic gene expression library

To provide a lytic proteome-wide panel of target antigens, we cloned a complete library of the known EBV lytic genes in expression vectors linked to *his* or *GFP* sequence tags. Supplementary Table 1 shows the list of lytic genes studied, their temporal phase within the lytic cycle, and the size and (where known) the function of the relevant protein product. Overall the library encompassed 70 genes encoding 2 IE, *32* E and *36* L proteins. Note that two genes, BNLF2a and BNRF1, were expressed as separate N-and C-terminal fragments, so that there were 72 constructs in all. In each case expression of the tagged protein was confirmed by immunoblotting (for *his*-tagged constructs) or by FACS analysis (for *GFP*-tagged constructs) of transiently transfected cells.

**Table 1:** HLA-A, B,C type of IM and HC donors. <u>*Footnote*</u>: HLA nomenclature in accordance with the IPD-IMGT/HLA database. Absence of an allele in homozygous individuals is represented by a dash. * IM269* was studied in acute IM and 4 years later as HC7*; note that another donor, HC2, happened to have the same HLA type as IM269/HC7 but the two individuals were not familiarly related.

| ID | HLA-A alleles |  | HLA-B alleles |  | HLA-C alleles |  |
| --- | --- | --- | --- | --- | --- | --- |
| IM84 | 02.01 | - | 15.01 | 44.02 | 03.04 | 05.01 |
| IM217 | 01.01 | 03.01 | 07.02 | 08.01 | 07.01 | 07.02 |
| IM223 | 02.01 | 24.02 | 08.01 | 40.01 | 03.04 | 07.01 |
| IM239 | 01.01 | 02.01 | 08.01 | 51.01 | 07.01 | - |
| IM243 | 01.01 | 23.01 | 08.01 | 44.03 | 04.01 | 07.01 |
| IM249 | 01.01 | 02.01 | 08.01 | 55.01 | 03.03 | 07.01 |
| IM269* | 03.01 | 11.01 | 15.01 | 35.01 | 03.03 | 04.01 |
| HC1 | 02.01 | 11.01 | 35.01 | 44.02 | 04.01 | 05.01 |
| HC2 | 03.01 | 11.01 | 15.01 | 35.01 | 03.03 | 04.01 |
| HC3 | 01.01 | 11.01 | 07.02 | 35.01 | 04.01 | 07.02 |
| HC4 | 01.01 | 02.01 | 39.01 | 40.01 | 03.01 | 12.01 |
| HC5 | 02.01 | 24.02 | 39.01 | - | 06.02 | 07.02 |
| HC6 | 01.01 | 02.01 | 44.02 | 57.03 | 05.01 | 07.01 |
| HC7* | 03.01 | 11.01 | 15.01 | 35.01 | 03.03 | 04.01 |

### T cell responses in IM patients

We used the above library to identify lytic antigen targets for the primary CD8+ T cell response to EBV in 7 acute IM patients who were serologically confirmed to be undergoing primary EBV infection. Table 1 shows the HLA-class I types of those 7 patients; collectively, they cover a range of 6 HLA-A alleles, 9 HLA-B alleles and 6 HLA-C alleles. Each of these alleles was individually cloned in expression vectors. Figure 1 illustrates the screening protocol used in the case of IM donors. The peripheral blood mononuclear cell (PBMC) population in IM is dominated by activated CD8+ T cells and, as in earlier work[12, 13], we directly expanded these activated cells by mitogen stimulation in IL2-conditioned medium. After 14 days CD4+ T cells were depleted, providing a polyclonal CD8+ effector population that could be used both for immediate testing and for cryostorage/re-testing. These effectors were screened on an EBV-negative antigen-presenting background, the MJS (Mel JuSol) melanoma-derived cell line[17], into which each of the 72 lytic gene constructs (and a GFP-tagged vector control) were individually introduced in combination with the donor’s two HLA-A allele or HLA-B allele or HLA-C allele constructs. Positive T cell recognition of an individual EBV antigen restricted through an HLA-A or −B or −C allele was detected by IFNg release and capture on a multi-well ELISA plate. Note that, where responses were identified, assays were subsequently repeated with individual HLA-expression constructs co-transfected with the relevant antigen in a second EBV-negative antigen-presenting background, the COS7 cell line[18]. This allowed us both to confirm the reproducibility of the antigen recognition and to identify the individual restricting element.

**Figure 1 :**
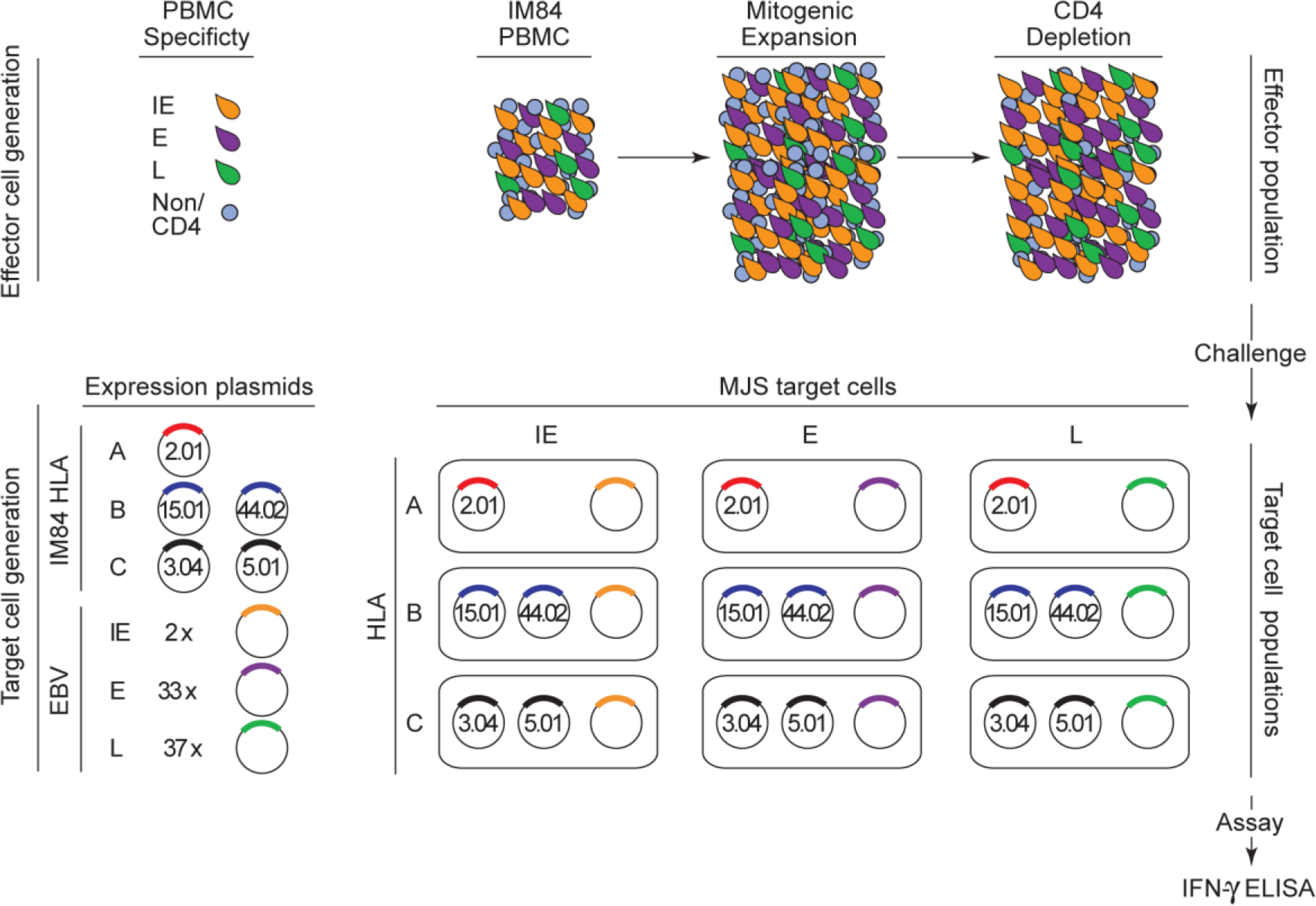
Schematic of the experimental design for proteome-wide screening of the primary CD8+ T cell response to EBV lytic cycle antigens in IM patients. The blood picture in acute IM is dominated by activated CD8+ T cells, the majority of which are EBV-specific and mostly directed against lytic cycle antigens; such cells are represented as activated lymphoblasts with IE (yellow), E (purple) or L (green) antigen specificity. Non-activated lymphocytes, including CD4+ T cells, are shown in blue. The activated cell population is preferentially expanded in vitro for two weeks in mitogen/IL2-containing medium, then depleted of any residual CD4+ T cells before testing as an effector population. Target cell populations, seeded in a multi-well format, are MJS cells transiently transfected with constructs expressing the IM donor’s two HLA-A alleles, two HLA-B alleles or two HLA-C alleles in combination with one of the 2 IE, 33 E or 37 L lytic gene constructs (or with a control construct); note that IM84 (HLA-A*0201, A*0201, B*1501, B*4402, C*0304, C*0501) is used as an example in the diagram. HLA-A, B or C-restricted recognition of a particular lytic gene construct is detected by IFNgamma release as measured in an ELISA assay.

Figure 2 shows results from the initial screening of one such acute patient, IM84. Targets are arranged vertically in blocs of IE, E and L proteins as in Supplementary Table 1 and, for each target, the levels of response through HLA-A, B and C alleles are shown in different colours. Positive responses were identified as those where IFNg release was at least >1.7-fold over background levels (i.e. that seen against a control construct) in the initial screen and reproducible in repeat assays. By these criteria, IM 84 effector population contained 25 individual responses, and the target range included 2/2 IE proteins, 12/32 E proteins and 7/36 L proteins. Based on levels of IFNg release, the most prominent responses were seen against the two IE proteins (BZLF1, BRLF1), four E proteins (BaRF1, BBLF2/3, BMLF1, BMRF1) and one L protein (BcLF1). IM84 had been chosen for analysis because this was the one individual from our earlier studies from whom cryopreserved acute phase cells were still available. In that earlier work, based on T cell cloning and testing on a limited number of targets, only one reactivity (against BMRF1) had been detected in IM84 [12]. The sheer range of reactivities identified in the present work bears witness to the value of proteome-wide screening and emphasises the multiclonal nature of the virus-induced response.

**Figure 2 :**
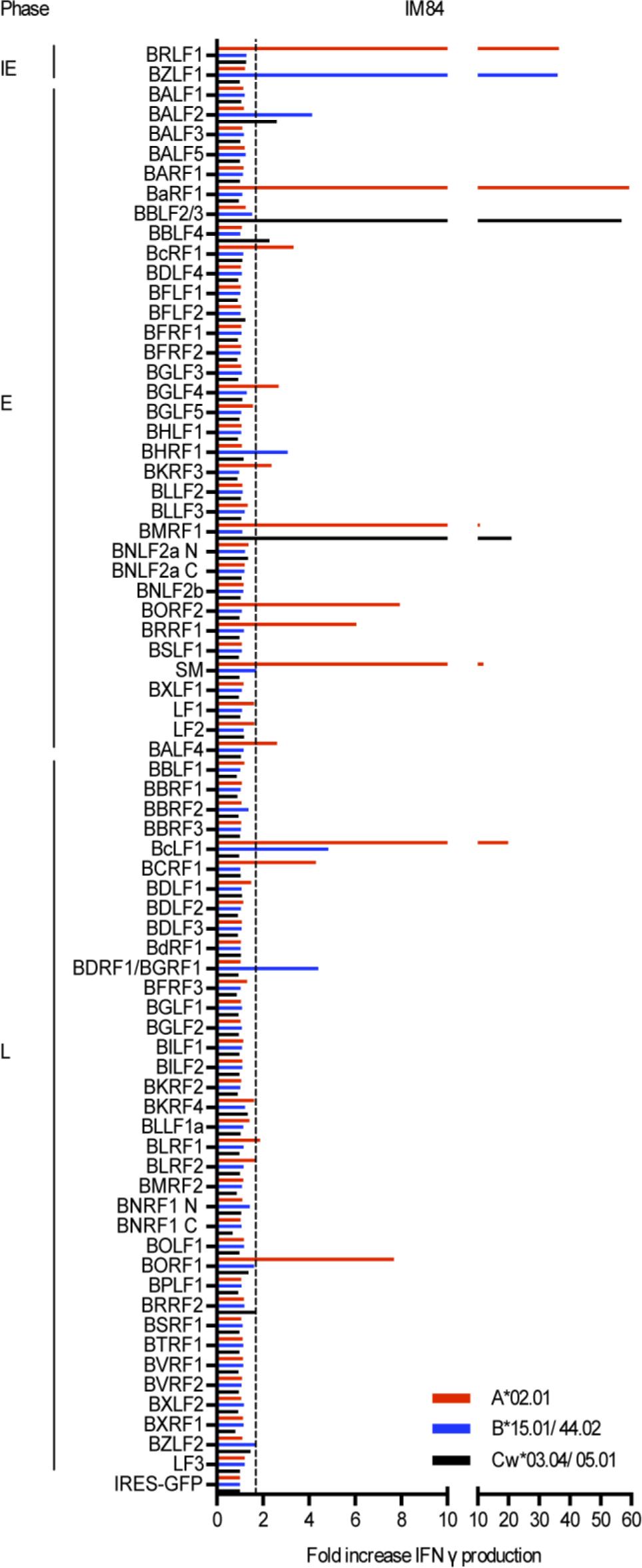
Results from proteome-wide screening of the primary CD8+ T cell response in IM 84 (HLA-A,B,C type as shown). The EBV lytic cycle targets are aligned vertically in blocs of IE, E and L antigens and responses are shown as fold-increases in IFNg production over the IRES-GFP control vector background; the dotted line indicates the cut-off identifying a positive response. For each target antigen, HLA-A, B-and C-restricted responses are shown separately as red (HLA-A), blue (HLA-B) and black (HLA-C) lines.

Detailed results from the other 6 IM patients are presented in Supplementary Figure 1 using the same format. The data again reveal the breadth of the primary response, with the number of distinct reactivities per individual patient being 15 (for IM217), 42 (for IM223), 5 (for IM239), 10 (for IM243), 35 (for IM249) and 14 (for IM 269). Figure 3 combines these individual results and provides an overview of the data from all 7 IM patients; where more than one subject responds to the same antigen/HLA A, B or C combination, the individual responses are marked by circles on the relevant line. Based on the observed levels of IFNg release, primary responses appear principally focused on the IE transcriptional activators, BZLF1 and BRLF1, and on certain E antigens, with prominent responses in several individuals against the ribonucleotide reductase subunits BORF2 and BaRF1, the major ssDNA-binding protein BALF2, the primase factor BBLF2/3 and the transcriptional activator/DNA polymerase processivity factor BMRF1; most of the above represent newly identified E antigen targets. L antigen-specific responses were less common by comparison but were clearly present in 6 of the 7 patients studied. Most of these L antigen-specific responses were relatively weak and, on the occasions where they were stronger, the target antigen tended to be seen by just one individual patient; interestingly, the best example of an L antigen being well recognised in more than one case involved the major viral capsid protein BcLF1. In total the data from all 7 IM patients showed 21 responses against the two IE proteins, 80 responses targeting 18 of the 32 E proteins, and 45 responses targeting 22 of the 36 L proteins. Overall, therefore, the proteome-wide screening of the primary CD8+ T cell response in IM revealed a much richer picture than that hitherto reported, with many new antigen-specific reactivities detected (see later Table); however, the same phase-dependent hierarchy of immunodominance among lytic cycle proteins observed in previous more limited studies (IE > E > L) [12] was still apparent.

**Figure 3 :**
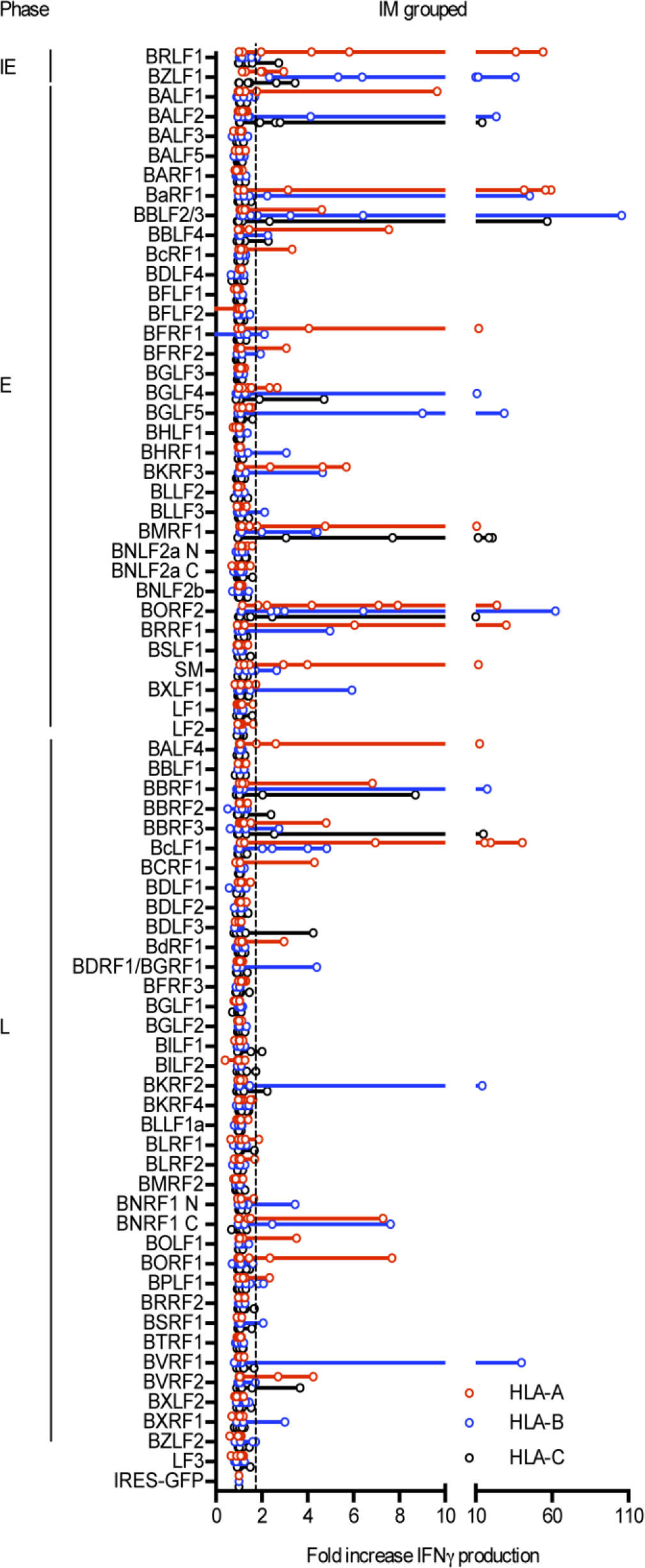
Combined results from proteome-wide screening of the primary CD8+ T cell responses in all 7 IM patients studied. Data are presented as in Figure 2, with individual IM patients’ results identified as open circles on each line.

### T cell responses in healthy virus carriers

We then sought to use the same proteome-wide screening to analyse lytic antigen-specific CD8+T cell memory in long-term virus carriers. This required a means of reactivating memory responses in vitro both efficiently and in a way that captured IE, E and L responses equally well. We reasoned that the conventional approach, stimulating PBMCs with the autologous EBV-transformed B lymphoblastoid cell line (LCL), had two disadvantages in that regard. Firstly, even in the most permissive LCLs, only a small fraction of cells enter lytic cycle; secondly, even where lytically-infected cells are present, EBV’s array of immune evasins expressed as early or late phase proteins[19–25] increasingly retard the presentation of later-expressed antigens on the LCL cell surface. We therefore adopted an alternative protocol, adapted from studies in the HSV system[3], which avoids the phase-specific effects of evasins and renders all lytic cycle antigens available for HLA I-mediated epitope display through cross-presentation in monocyte-derived dendritic cells.

The experimental protocol is illustrated in Figure 4. From each healthy carrier studied, dendritic cells were prepared by IL4/GMCSF-stimulation of CD14+monocytes as described [3]. These cells were then exposed for 16-20 hrs to a lysate of the HEK-293 epithelial cell line (American Type Culture Collection) carrying the recombinant EBV strain B95.8 genome, made 48 hrs after triggering the latter’s induction into lytic cycle, a time judged to be optimal to capture the full gamut of lytic cycle proteins[26]. The antigen-loaded dendritic cell preparation was then co-cultured with autologous lymphocytes and, as described [3], antigen-stimulated memory cells recognised through cell surface up-regulation of the CD137 activation marker. Co-cultures were harvested on day 3 and, following cell surface staining, the CD137+, CD8+ fraction was sorted by FACS and then expanded for 2 weeks in mitogen/IL2-conditioned medium just as had been used to expand the IM effectors. The resultant effector population was then screened on the entire EBV lytic antigen panel using the same combinations of antigen/self HLA-A, B and C alleles and multi-well readout as in the IM work above.

**Figure 4 :**
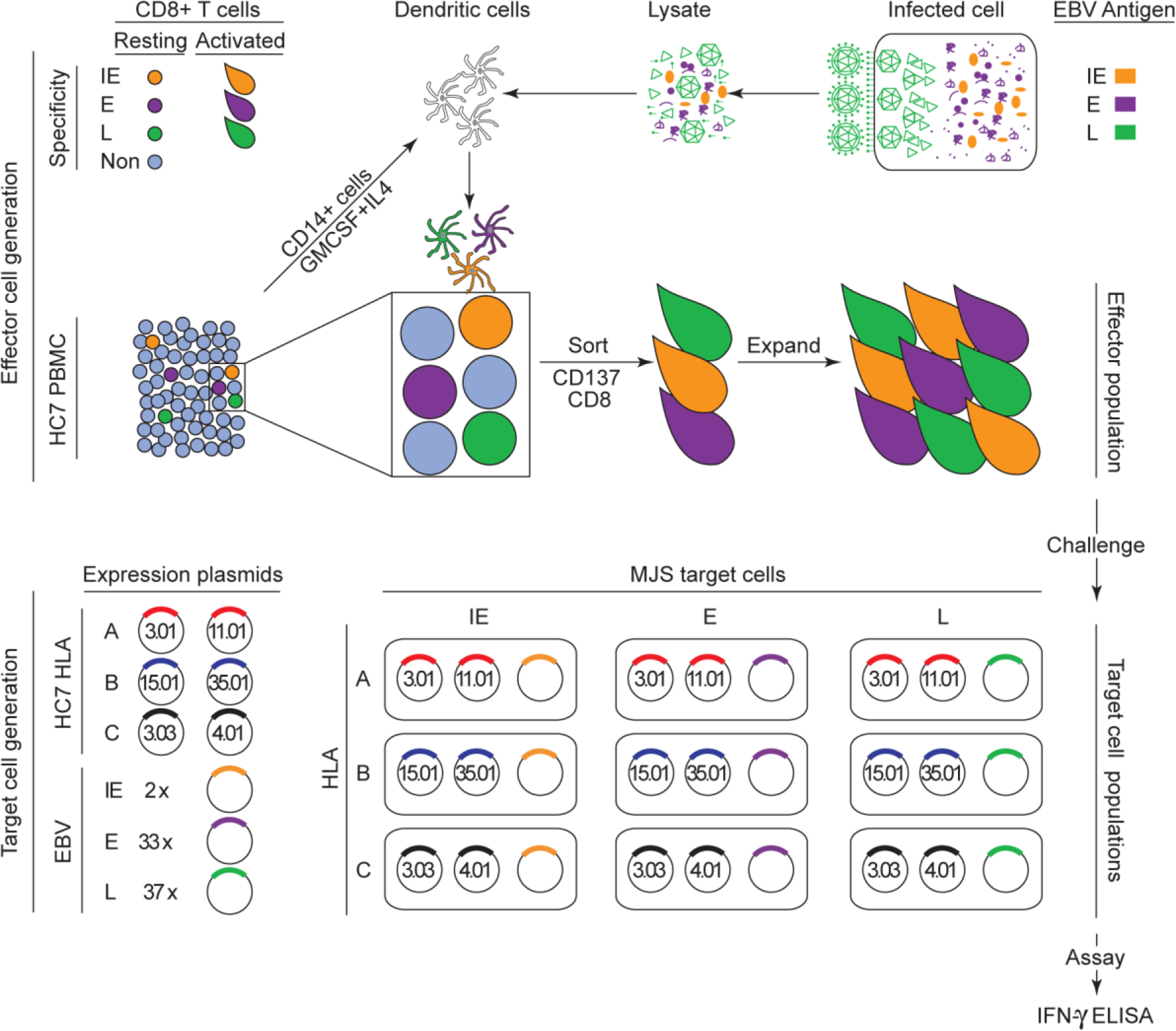
Schematic of the experimental design for proteome-wide screening of the memory CD8+ T cell response to EBV lytic cycle antigens in healthy virus carriers. EBV-specific CD8+ memory cells in the blood of healthy carriers constitute a small fraction of the resting lymphocyte pool and require selective re-activation before expansion in vitro. To achieve this, dendritic cells are first prepared from the healthy carrier by GMCSF/IL4 treatment of CD14+ blood monocytes and then exposed to an IE, E and L antigen-containing lysate made from EBV lytically-infected 293 cells. Lysate-loaded dendritic cells displaying cross-presented lytic antigens are then co-cultured with PBMCs of the healthy carrier, leading to re-activation of the EBV lytic antigen-specific memory population. After 3 days, CD8+ T cells now expressing the CD137 activation marker are then selected by CD8+/CD137+ sorting. Thereafter the sorted cells are expanded in vitro for two weeks in mitogen/IL2-containing medium (as used for IM cell expansion) and tested as an effector population. Testing is carried out on transiently transfected MJS cells using the relevant HLA/lytic gene combinations as described in Figure 1; note that HC7 (HLA-A*0301, A*1101, B*1501, B*3501, C*0303, C*0401) is used as an example in the diagram.

As a first step, we took advantage of the fact that one of the IM patients studied (IM269) was available for sampling four years later, long after resolution of primary infection (HC7, post-IM269). Figure 5 presents the results of target antigen mapping at this time point alongside that from the original acute IM bleed. While the acute IM results had shown the typical focus on IE and E targets, the overall picture 4 years later had a different emphasis. Several of the original reactivities (for example to the IE antigen BZLF1 and to the E antigens BGLF5, BKRF3 and BORF2) had been retained but were generally less prominent, while, as noted in earlier work [ref 13], other E antigen responses had fallen below the level of detectability post-IM. By contrast, an initially weak response to the L antigen (BcLF1) had become the most prominent and had been joined by strong responses to another L antigen BNRF1.

**Figure 5:**
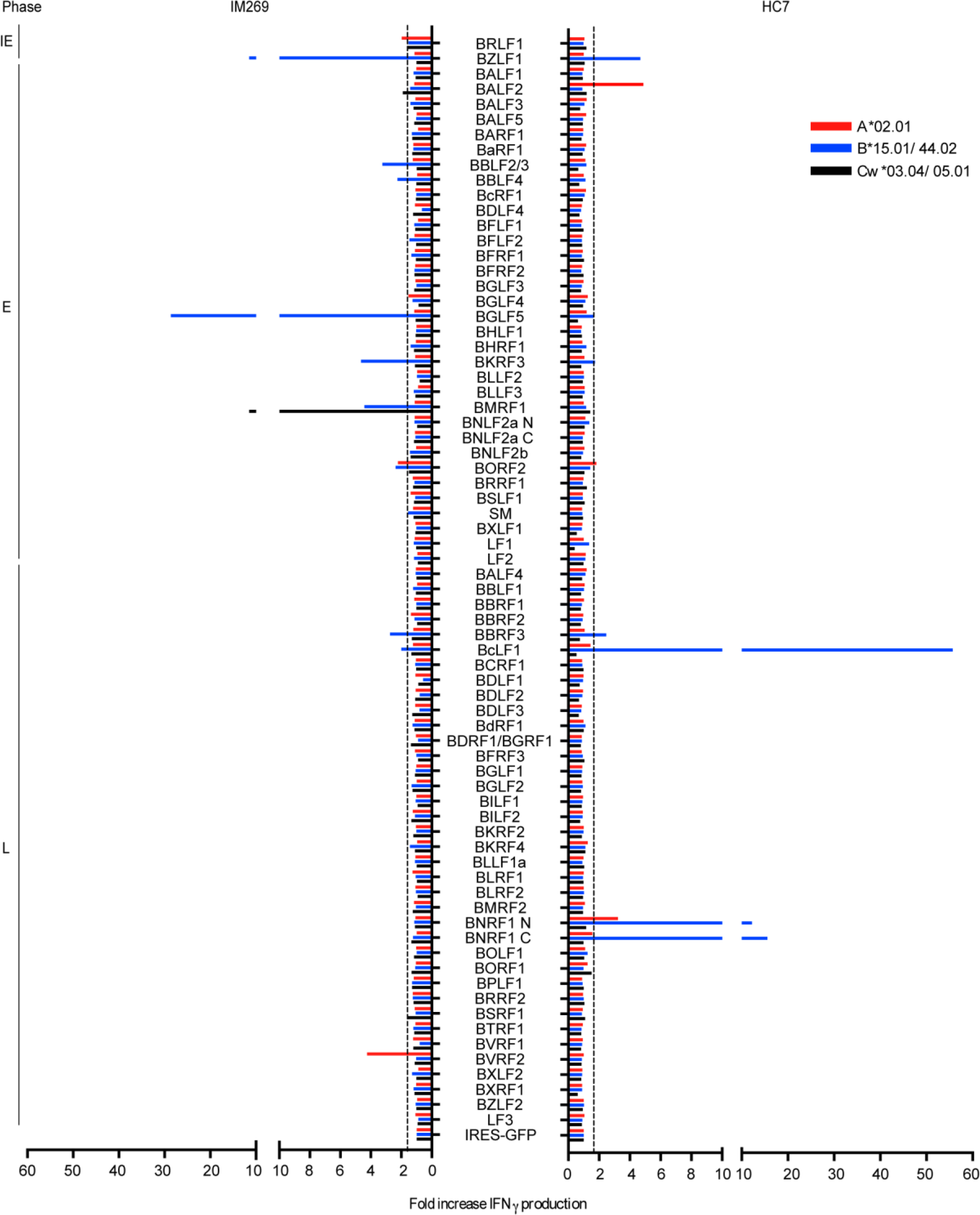
Results from proteome-wide screening of the primary and memory CD8+ T cell responses in a single individual. Donor IM 269 was screened in the acute phase of IM and, 4 years later, as donor HC7; the individual’s HLA-A,B,C type is as shown. Data are presented as in Figure 2.

In all we studied 7 healthy long-term virus carriers in this way. They are listed in Table 1 along with their HLA I types; collectively they cover a range of 5 HLA-A alleles, 7 HLA-B alleles and 8 HLA-C alleles, many of which were also present in the IM patient panel. Again, each of these alleles was individually cloned in expression vectors as above to allow definitive restriction of responses. Figure 6 shows the results from one such healthy carrier, HC1. The overall picture is broadly similar to that seen for HC7 above; in this case 15 responses were detected and they were distributed across the IE, E and L antigen spectrum. Thus prominent responses to the IE antigens BRLF1 and BZLF1 and to four E antigens (BaRF1, BMRF1, BORF2 and SM) were accompanied by equally prominent responses to four L antigens (BBRF3, BcLF1, BDLF1 and BNRF1). In the case of HC1, subsequent mapping of these responses to individual HLA alleles (as described later) identified four of the above IE or E antigen-specific responses where the major peptide epitope presented by these alleles, namely A*0201-or B*3501, was already known. This allowed us to conduct an internal control, using an IFNgamma-based Elispot assay to measure the memory responses to these four peptide epitopes as present in donor HC1 PBMCs ex vivo. The data (see Figure 6 legend) show that the size order of those ex vivo responses had indeed been retained in the in vitro-reactivated and expanded effectors used in this work. Further results from proteome-wide screening of in vitro-expanded memory preparations from the other 5 healthy carriers are shown in Supplementary Figure 2, with the number of responses seen per donor ranging from 19 (in HC2), 25 (in HC3), 9 (in HC4) and 16 (in HC6) down to just a single response reproducibly observed in HC5. In each case the overall distribution of target antigen choice looked different from that typically seen in IM.

**Figure 6 :**
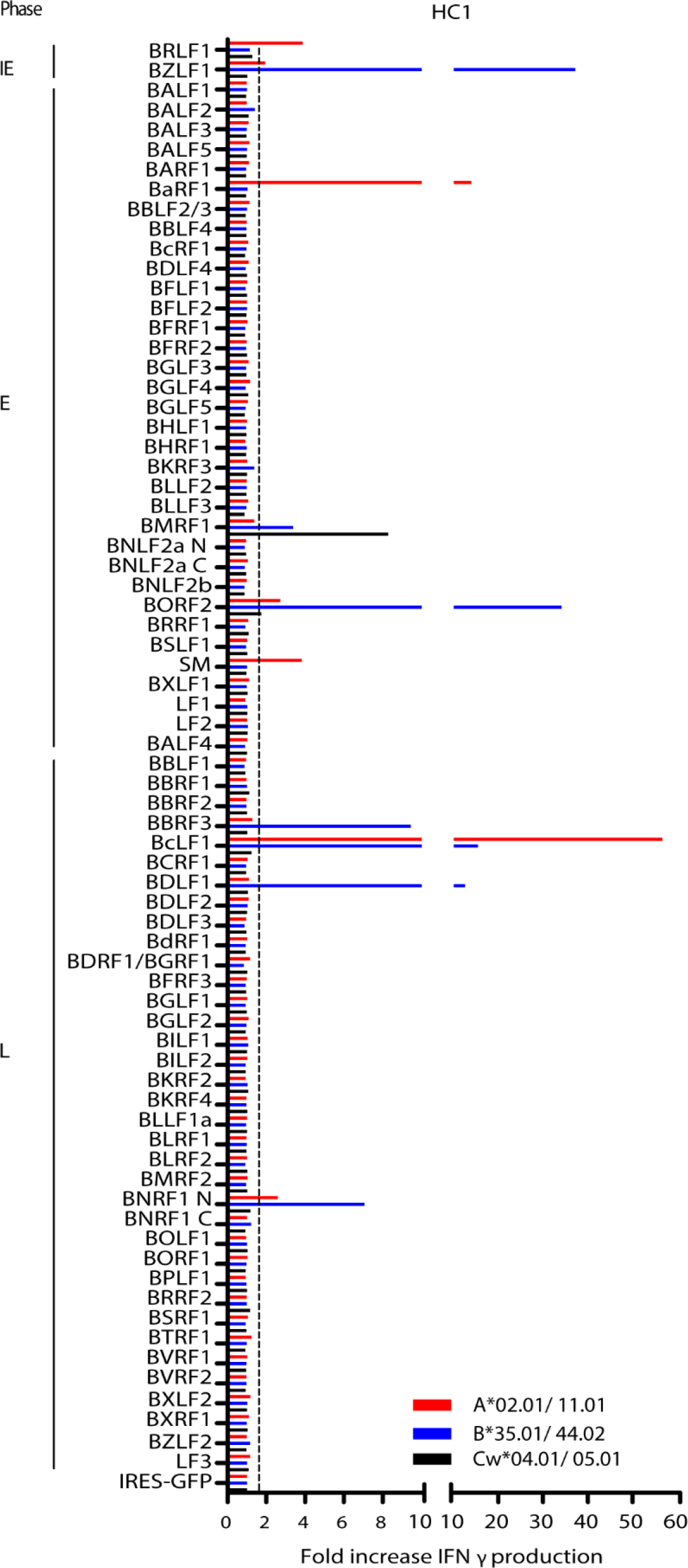
Results from proteome-wide screening of the memory CD8+ T cell response in HC1 (HLA-A,B,C type as shown). Data are presented as in Figure 2. Of the 15 responses detected in memory cell preparations from this donor, there were 4 responses which (following restriction to individual HLA alleles) aligned to previously defined IE or E antigen/HLA target combinations for which the major peptide epitope was known[8]; these were BZLF1/HLA-B*3501 (peptide EPLPQGQLTAY), BaRF1/HLA-A*0201 (peptide LLIEGIFFI) BRLF1/HLA-A*0201 (peptide YVLDHLIVV), and SM/HLA-A*0201 (peptide GLCTLVAML). This allowed us to compare the size order of these responses in in vitro-expanded HC1 effectors as shown in the Figure (fold-increases in IFN-γ release of 37.2, 14.0, 3.9 and 3.8 respectively) with the size order seen when PBMCs from the same donor were tested in an ex vivo IFN-γ-based Elispot assay against the above peptides (spot-forming cell numbers per 106 PBMCs of 1228, 1148, 520 and 358 respectively). The same order was observed in the two populations, suggesting that in vitro activation and expansion gives a fair representation of the content of in vivo memory.

Figure 7 presents the combined results from all 7 healthy carriers (using the same format as for the IM summary data shown earlier). This illustrates the distinct character of the memory response. While the two IE proteins and certain early proteins such as BALF2, BaRF1, BMRF1 and BORF2 remain frequently preferred targets, L antigen-specific reactivities are now prominent components of the memory response with particular target antigens, notably BBRF3, BcLF1, BDLF1 and BNRF1, often more prominent than IE and E targets. Overall the healthy carriers mounted 8 responses against the 2 IE proteins, 37 responses targeting just 14 of the 32 E proteins, and 51 responses targeting 15 of the 36 L proteins. This distribution, with > 50% of detected healthy carrier responses against L antigen targets, is clearly different from that seen in IM where only 30% responses were so directed. The contrast is even more marked when one compares the relative prominence of the IE, E and L responses in the two situations. As shown in Figure 8, while individual responses to the different classes of target antigen occupy a wide range in both donor cohorts, IE and E-specific responses tend to be stronger in IM while L-antigen-specific responses tend to be the strongest in healthy carriers.

**Figure 7:**
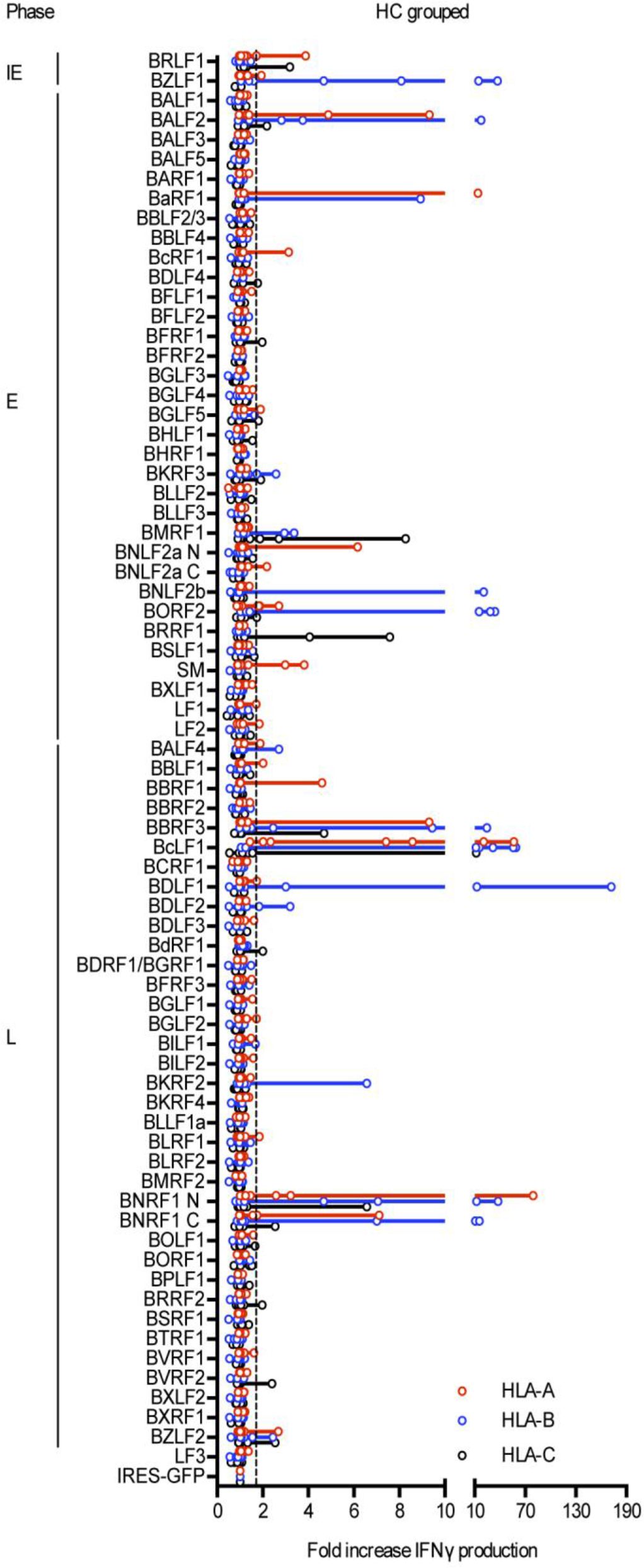
Combined results from proteome-wide screening of the memory CD8+ T cell responses in all 7 healthy carriers studied. Data are presented as in Figure 2, with individual results identified as open circles on each line.

**Figure 8:**
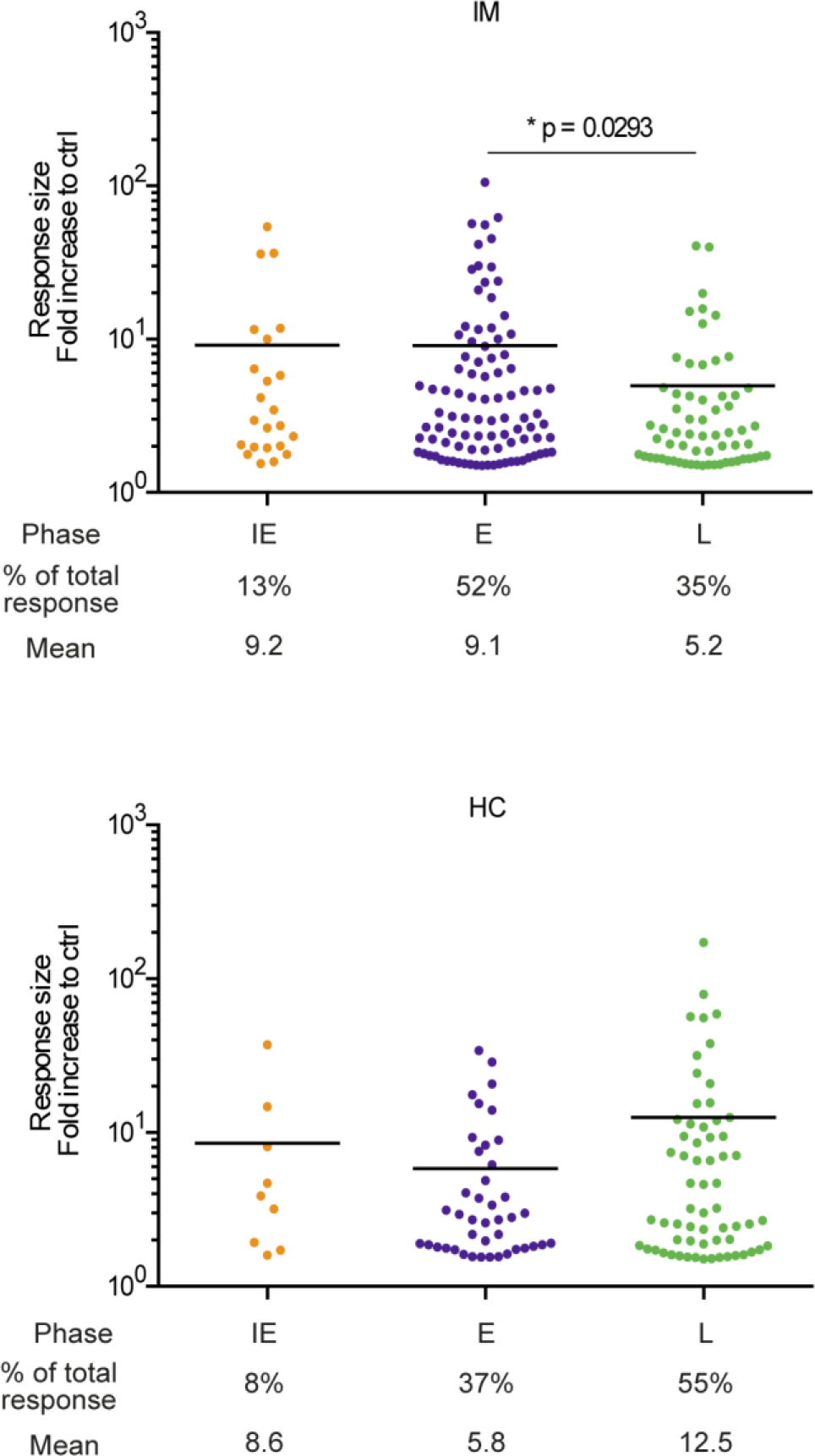
Comparison of individual response sizes to IE (orange), E (purple) and L (green) antigens detected in the 7 IM patients (top graph) and 7 HC donors (bottom graph). Shown below the dot plots are the number of responses to IE, E and L antigens as a proportion of all responses, and the mean size of those responses; response size is defined as the fold-increase in IFNg production over the GFP control vector background In IM patients, IE and E responses are on average larger than L responses (E is significantly larger than L, p=0.0293) and 52% responses are directed against E antigens: in HC, L responses are on average larger than IE and E responses and 55% responses are directed against L antigens.

As a prelude to further analysis of the data, Figure 9 presents the individual IM patient and healthy carrier responses to the full spectrum of lytic cycle antigens in checkerboard format, with three vertical columns per individual distinguishing reactivities that are restricted through one of the donor’s two HLA-A, −B or −C alleles. Positive responses are marked by shaded squares, with red squares identifying the strongest responses (>5-fold increase in IFNg release over background).With this as a reference point, we set out to address two further questions.

**Figure 9:**
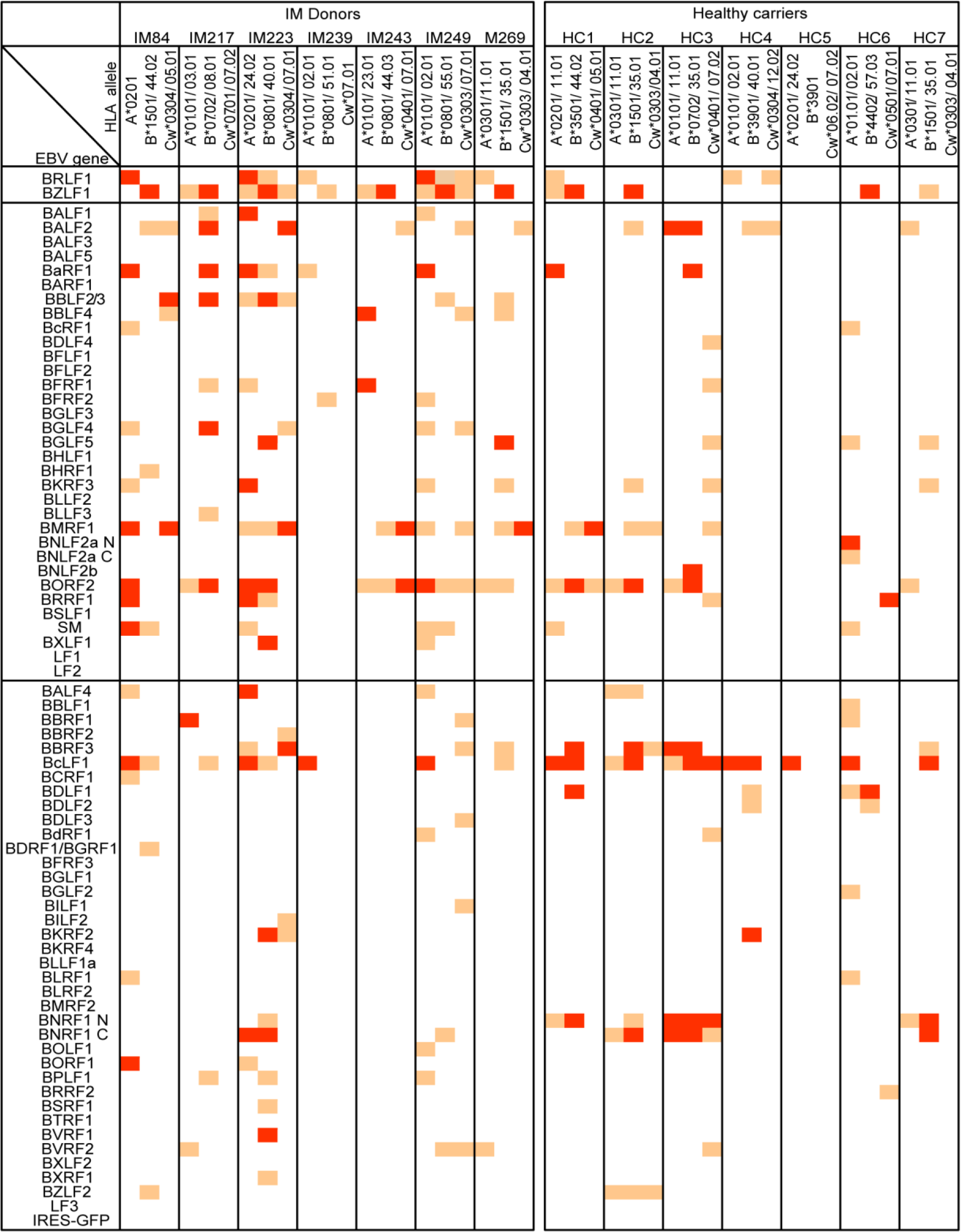
Summary of all individual responses detected in the 7 IM patients and 7 healthy carriers studied. Lytic gene targets are arranged vertically in blocs of IE, E and L antigens as in Figure 2. Individual responses mapping to the donor’s HLA-A, B or −C restricting alleles are identified as orange and red squares; red squares denote the stronger responses (IFNg release >5-fold greater than background).

### Immunogenic EBV antigen/HLA allele combinations and prevalent responses

In a subsequent set of experiments, effector cell preparations from all the above IM patients and healthy carriers were re-screened on EBV-negative COS7 cells transiently transfected to express the target antigen of interest and one of the two candidate HLA alleles. This formally identified the antigen/HLA I restricting allele combinations against which responses were directed. Supplementary Table 2 lists the antigens presented through the individual HLA-A, HLA-B and HLA-C alleles, and the donors (IM or HC) who made a relevant response. These include (i) 51 antigen-specific responses restricted through one of 6 HLA-A alleles that are common or relatively common in Caucasian populations, namely HLA-A*0101, A*0201, A*0301, A*1101, A*2301 and A*2402, (ii) 73 responses restricted through one of 9 HLA-B alleles, including common alleles such as HLA-B*0702, B*0801, B*1501, B*3501 and B*4001, and (iii) 31 responses restricted through one of 7 HLA-C alleles. Based on existing records of known EBV-specific CD8+ T cell responses [8] (the Immune Epitope Database IEDB.org), a large majority of these antigen/HLA allele target combinations are novel. Among the most noteworthy are the 6 antigens found to elicit an HLA-A*0101-restricted response, since these represent the first well documented examples of this common HLA allele mediating a response against any EBV antigen, lytic or latent. Likewise the number of HLA-C-restricted lytic cycle responses is unexpectedly high when compared to latent antigen responses which rarely if ever display HLA-C restriction.

We went on to ask whether the above data allowed one to identify antigen/HLA I combinations that induced “prevalent” responses, i.e. where a response to that combination was seen in the majority of individuals possessing that HLA I allele[27]. Such combinations are shaded in Supplementary Table 2, alongside the number of relevant donors studied (ranging between 2 and 8 per allele). Prevalent responses were seen in the context of three HLA-A alleles (A*0201, A*1101, A*2402), six B alleles (B*0702, B*0801, B*1501, B*3501, B*4001, B*4402) and three C alleles (C*0303, C*0304, C*0401). Many of these are again novel. Interestingly, there were also several instances where prevalent responses on different HLA I backgrounds were directed against the same viral antigen. These target antigens were not drawn from one phase of the cycle, however, but straddled across the IE/E/L range. For example, BZLF1 (IE phase, transactivator) was a prevalent target on four alleles and a target on 11 alleles in total, BORF2 (E phase, ribonucleotide reductase large sub-unit) was a prevalent target on three alleles and a target on 11 alleles in total, and BcLF1 (L phase, major capsid protein) was a prevalent target on five alleles and a target on 9 alleles in total.

Identifying the HLA restricting alleles allowed us to determine the relative number of HLA-A-versus B-versus C-restricted responses seen in IM versus healthy carriers but also the relative strength of those responses in the two situations. The data are shown in Figure 10. Compiling the results from all 7 IM patients, HLA-A-and HLA-B-restricted responses were both more common than those through HLA-C (60 and 56 versus 30 responses respectively) and on average slightly larger. By contrast, analysing the healthy carrier results in the same way, HLA-B-restricted responses were not only the most common (41 versus 35 HLA-A-and 20 HLA-C-restricted) but also gave significantly stronger responses than both their HLA-A-and C-counterparts (HLA-B versus A: P<0.01, HLA-B versus C: P<0.001. Kruskal-Wallis test with a Dunn’s multiple comparison). Notably the greater contribution from HLA-B alleles in virus carriage was not just a consequence of the shift towards L antigen targets, since the same trend was seen comparing the HLA restriction of IM and carrier responses against the separate blocks of IE, E and L antigens.

**Figure 10:**
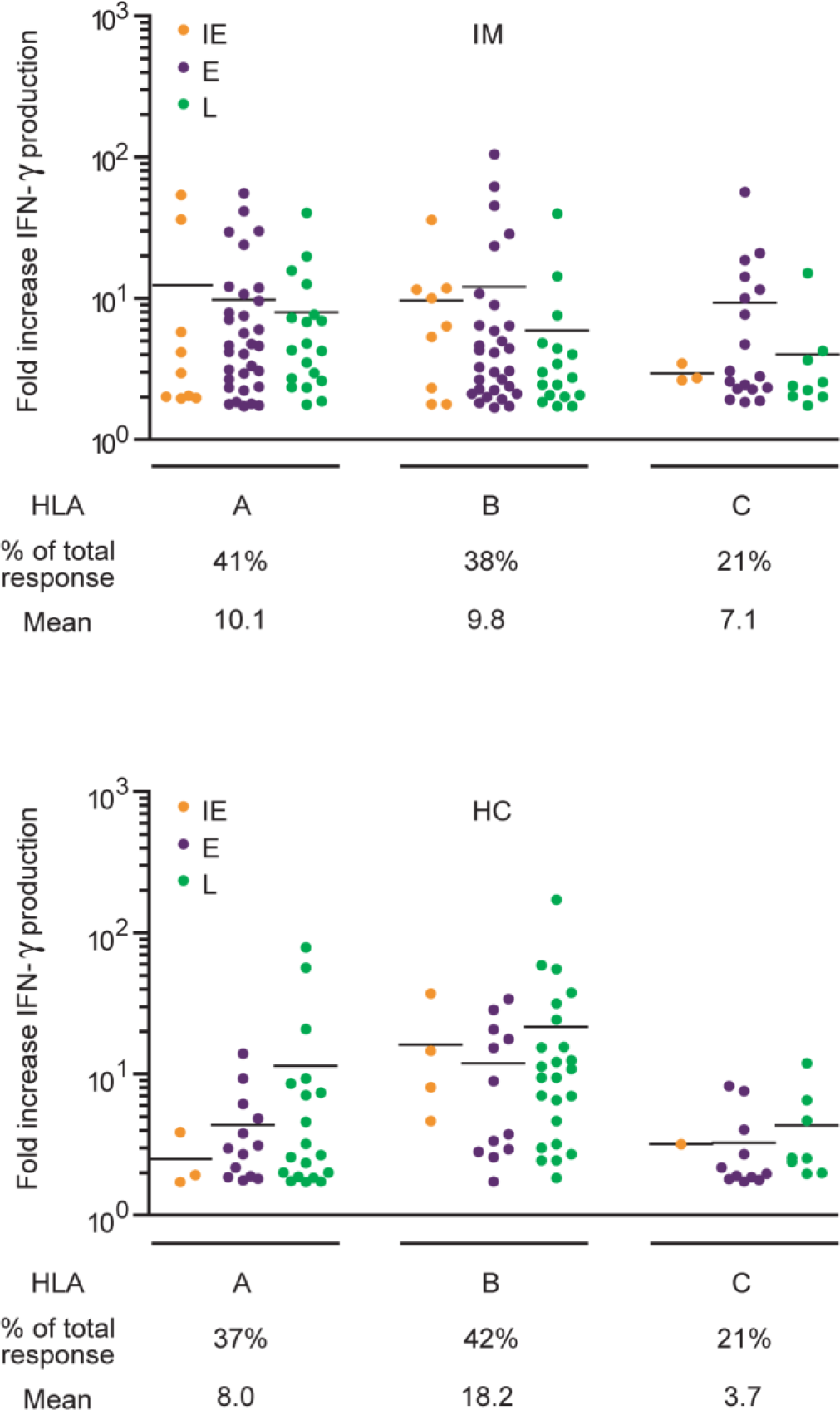
Comparison of individual response sizes restricted through HLA-A versus HLA-B versus HLA-C alleles in the 7 IM patients (top graph) and 7 HC (bottom graph). Shown below the dot plots are the number of responses restricted through those alleles as a proportion of all responses, and the mean size of those responses. Colours indicate individual responses to IE (orange), E (purple) and L (green) antigens. In IM, HLA-A and B-restricted responses are more common than those restricted through HLA-C and have a slightly higher mean; by contrast in healthy carriers, HLA-B-restricted responses are the most common and are larger than both HLA-A-restricted and HLA-C-restricted responses.

### Antigen size in relation to response strength

As the above work has shown, CD8+ T cell responses to EBV are most often complex mixtures, with up to 42 distinct reactivities detectable in any one individual. The variety of HLA types among both IM patients and healthy carriers would be expected to diversify target choice across the range of lytic cycle proteins. However a subset of proteins appear to preferentially attract responses, based on the frequency with which they are detected as targets (orange shading) and especially as the most prominent targets (red shading) in Fig 9. Moreover the hierarchy of responses to individual proteins is different in IM versus healthy carriers. To provide a measure of antigenicity for each target detected, we summed all of the response sizes seen to that antigen in the interferon-gamma assays in IM patients and, separately, in healthy carriers. In each case we then plotted the cumulative size of the response against the size of the antigen (expressed as number of unique amino acid residues). We reasoned that, if all viral proteins were equally available for presentation to the CD8+ T cell repertoire in vivo, then one might anticipate that the larger proteins (i.e. proteins containing a greater number of unique peptides) would attract a correspondingly large proportion of overall responses. Interestingly, as shown in Fig 11, there was no obvious link between the two parameters when looking at the IM data, where strong responses against small-to-medium size IE and E antigens such as BZLF1, BaRF1, BMRF1 and BRLF1 were a major feature. In contrast response strength and antigen size were significantly correlated in healthy carrier responses, where the strongest cumulative responses were against large L antigens such a BcLF1 and BNRF1. Such a result is at least consistent the idea that antigen display on cells inducing the primary response is not a fair representation of the entire EBV lytic cycle proteome, whereas this bias is removed in the long-term carrier state.

**Figure 11:**
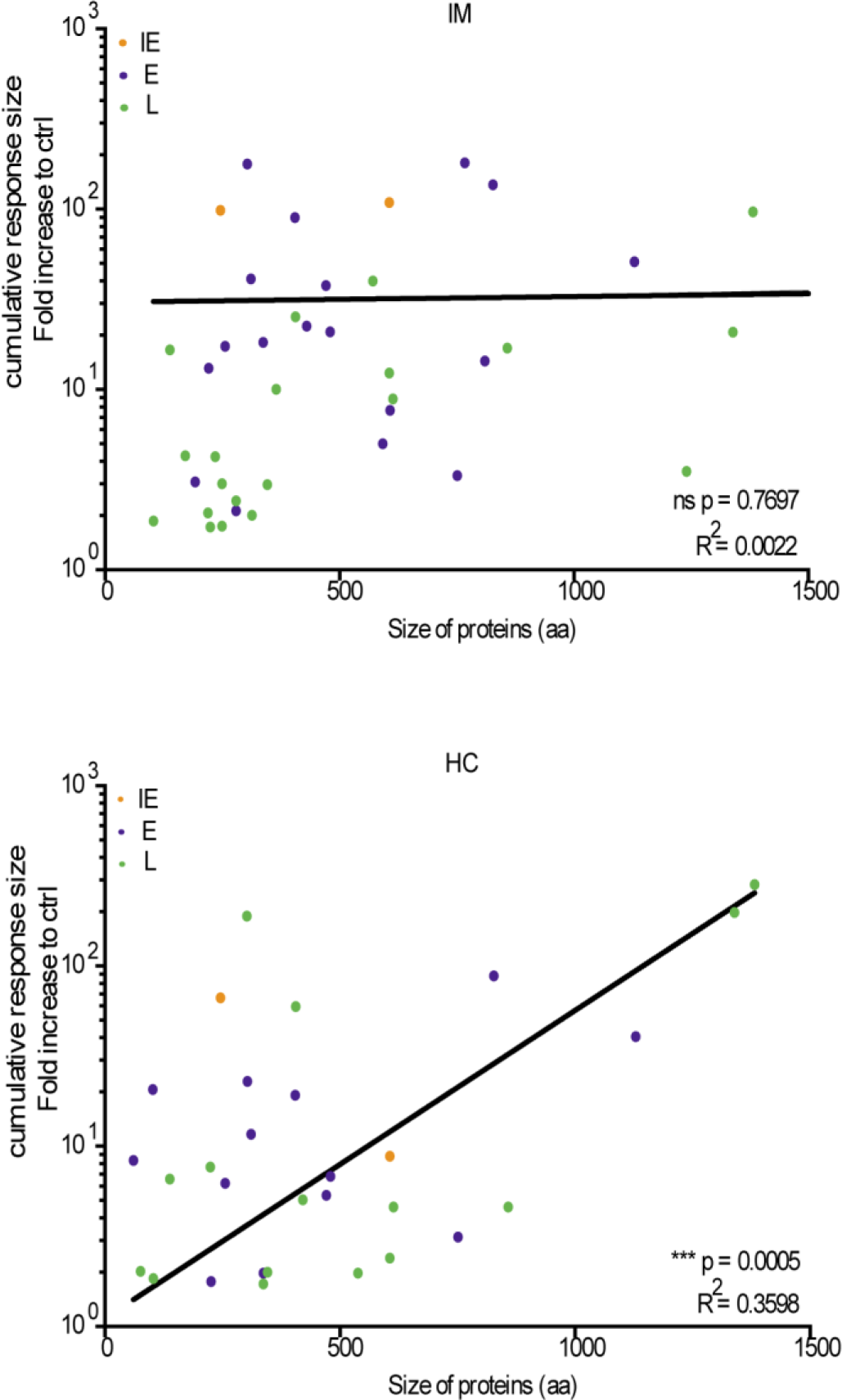
Comparison of the cumulative size of IM patient and healthy carrier responses to individual target antigens in relation to antigen size. For each group of donors, the response size shown for each antigen represents the sum of all individual response sizes to that antigen. Antigen size is shown as the number of amino acids in the primary sequence (allowing for any repeat sequences) as in Supplementary Table 1. Colour coding identifies IE (orange), E (purple) and L (green) antigens. Note that only those antigens that induced a detectable response are included in this analysis; for the purpose of presentation, we do not show the very small IM donor response to the large BPLF1 protein. Immunogenicity and antigen size were significantly correlated in healthy carrier responses (P=0.0005, R^2^=0.3598).

## DISCUSSION

Here we report the first comprehensive study of CD8+ T cell responses to EBV lytic cycle proteins and the first proteome-wide comparison of CD8 target antigen choice in primary versus persistent infection for any human herpesvirus. We began by screening effectors from 7 acute IM patients on the full panel of 70 EBV lytic cycle proteins. This provided a much richer picture of the primary CD8 response than seen hitherto, with a mean of 21 and a range of 5-42 antigen-specific reactivities detected per patient. While there are no comparative data from primary infections with other human herpesviruses, such breadth is broadly in line with that seen in mice responding to murine cytomegalovirus (MCMV) and murine gamma2-herpesvirus (MHV68) challenge [28–30]. The new data reinforced certain aspects of earlier work on IM, in particular the high incidence and strength of responses against the two IE transcriptional activators, BZLF1 and BRLF1, and to a lesser extent against two E proteins, BALF2 and BMRF1, with roles in viral DNA replication [12]. However, the data also identified three prominent new E antigen targets, BBLF2-3, the primase factor, and BORF2 and BaRF1, large and small sub-units of the viral ribonucleotide reductase. While L-specific responses were frequently present in IM, they were generally weak. Only one L antigen, the major capsid protein BcLF1, came close to the dominant IE and E targets in terms of the frequency and size of responses, and this was mainly attributable to the overall impact of an A*0201-restricted reactivity.

Parallel work on healthy carriers, including one individual also studied in primary infection (IM269/HC7), showed a relative shift in the distribution of responses compared to IM with L antigen targets gaining greater emphasis. Responses to some of the strongest IE and E antigen targets were still frequently present. However the most prominent response in every healthy carrier tested was directed against an L antigen, most often against one of three virion structural proteins, the major capsid protein BcLF1, the large tegument protein BNRF1 and the envelope glycoprotein BBRF3 (gM). The concept that virus carriage brings an increasing emphasis on L antigen targets also chimes with published data from rhesus macaques carrying the EBV-related rhesus gamma-1 herpesvirus, rh-LCV, where memory cell responses were equally split between the 10 E and 8L rh-LCV antigens tested, with a hint of greater L focus in older animals[15]

We considered the possibility that the above differences were artefacts of the different protocols used to generate primary and memory effectors, and/or by the process of in vitro expansion per se. However we do not believe this to be the case. Primary effector preparations were made by direct expansion of the activated CD8+ T cells that dominate the blood picture in IM, and earlier work using defined HLA/EBV peptide tetramers has shown that such ex vivo-expanded IM effectors accurately reflect the in vivo response[12]. By contrast, the small population of EBV-specific memory cells in the blood of healthy carriers need to be selectively reactivated in vitro before expansion; not surprisingly, the only published attempt at direct memory cell expansion ex vivo revealed very few detectable EBV-specific reactivities despite screening on a wide target panel[31]. To avoid the bias inherent in using LCL cells to reactivate memory, we adapted an approach first developed for HSV[3] whereby monocyte-derived dendritic cells cross-present the whole spectrum of virus-coded lytic antigens to the CD8 memory pool. We therefore induced latently-infected 293 cells into lytic cycle and made total cell extracts at a time when >50% cells had already entered late phase and which recent proteomic analysis suggests is optimal for the most complete representation of all lytic antigens[26]. While it is possible that some antigens were absent or under-represented in such extracts, the fact that memory responses were reactivated across the whole spectrum of IE, E and L antigens suggests that antigen representation is not a major problem. Indeed among the targets of these reactivated responses were antigens such as the immune evasin BNLF2a, which is only transiently detectable early in lytic cycle[20], and the portal and scaffold proteins BBRF1 and BdRF1, known to be the least abundant components of the capsid in virus particles[32] and reportedly present at very low concentrations in lytically-infected cells[26]. These considerations and the fact that, for IE and E antigens with already defined target epitopes, the relative strength of antigen-specific memory responses seen in expanded memory cell preparations broadly reflected that seen in ex vivo Elispot assays suggests that such preparations fairly represent the content of virus-specific memory.

In all but one case our IM and healthy carrier cohorts involved different individuals, but it seems unlikely that chance differences in HLA type between the two groups could account for the different patterns of responses. Of the 26 HLA I alleles covered by the present study, 16 alleles (5 HLA-A, 5 HLA-B and 6 HLA-C) are shared between the two donor groups; witness also the evidence from IM269/HC7 where a shift in target choice was seen over time in the same individual. Assigning responses to individual HLA alleles defined >150 different lytic antigen/HLA target combinations, many of which are novel, and identified a subset of those that were frequently recognised in individuals with the relevant allele. Interestingly HLA-A and HLA-B-restricted responses were equally matched in both incidence and size in primary infection, whereas HLA-B-restricted responses appeared to make the greater contribution in memory. A predominance of HLA-B-restricted responses has been observed in HIV[33], measles[34] and mycobacterial[35] infections and was indeed suggested in an earlier study of virus carriers using EBV epitope peptides[36]. It is also worth noting that, in both phases of infection, around 20% lytic antigen-specific responses were HLA-C-restricted. This contrasts sharply with EBV latent antigen-specific responses where examples of HLA-C restriction are extremely rare [8, 37]. Finally, one of the more interesting comparisons between individual alleles involved HLA-A*0101 and A*0201 because of their identification as high and low risk alleles respectively for the development of EBV-positive Hodgkin lymphoma [38–40], a disease potentially linked to impaired T cell surveillance[39, 41]. In that regard, previous studies have defined a small number of EBV latent and lytic epitopes restricted though A*0201, whereas numerous studies have failed to reveal any EBV-specific responses restricted through A*0101[8, 42]. The present work extends this analysis significantly; HLA-A*0201 was indeed the stronger restricting allele, but not overwhelming so. Overall A*0201-restricted responses were detected against 20 different lytic cycle antigens among the A*0201-positive donors studied (4 IM, 4 healthy carriers). By comparison, A*0101 restricted responses were seen against 6 different antigens among a similar number of A*0101-positive donors (4 IM, 3 healthy carriers); it may be significant that each of these novel A*0101-restricted responses was relatively weak and, with one exception, only detected in IM.

With respect to the data from IM patients, our proteome-wide screening greatly strengthens the evidence for an IE > E > L hierarchy among lytic cycle antigens as targets of the EBV-induced primary response. Note that this hierarchy parallels the efficiency with which IE, E and L antigens are presented on the surface of lytically-infected cells because, as cells move through lytic cycle, endogenous antigen presentation is progressively impaired by virally-coded evasins [19–24, 43] and so epitopes derived from earlier expressed antigens are preferentially displayed. This parallel leads us to suggest that the primary response to EBV may be largely driven by direct contact between the CD8+ T cell repertoire and infected cells themselves. Thus the two IE proteins, BZLF1 and BRLF1, are dominant targets and together attract ~15% of all IM responses while accounting for only ~3% of the total lytic protein sequence space. A further 55% of all IM responses are directed against E proteins, although it must be stressed that E antigen choice is not evenly distributed. There are large differences between individual E proteins, with almost half of the responses being directed against just 5 dominant targets (BALF2, BMRF1, BBLF2-3, BORF2 and BaRF1). It may be that the timing of these particular proteins’ expression renders them less protected by the early evasin BNLF2a and/or the early/late evasin BILF1. However this remains unclear because the precise chronology of antigen expression in early phase is poorly defined. Other factors, such as the rate of de novo antigen synthesis, may be at least as important and it is significant that BALF2, BMRF1, BORF2 and BaRF1 are among the most highly expressed E antigens in lytic cycle [26, 32].

Our parallel analysis of CD8+ T cell responses in healthy carriers revealed an increasing focus on L phase antigens. More than 50% of detectable memory responses were directed against L antigens compared to only 30% in IM. This shift leads us to speculate that, during years of virus carriage, recurrent entry of latently-infected B cells into lytic cycle provides a bolus of lytic antigens, including L antigens, that are cross-presented to CD8+ T cells and this switch in the main pathway of antigen presentation re-shapes the memory response. Again, however, there are large differences between the 36 L phase proteins, with three dominant targets (BcLF1, BNRF1 and BBRF3) attracting more than half of all L-specific responses. Note that of these, BcLF1, the major capsid protein and BNRF1, the major tegument protein, are both in the top 15 EBV proteins detected in lytically-infected cells by proteomic analysis[26] but, interestingly, are also the two most abundant proteins in the proteome of mature virus particles[32] ; BBRF3, the gM glycoprotein is reportedly much less abundant but, as a glycosylated protein, may have been under-scored in these proteomic assays. It therefore seems possible that, as well as lytically-infected cells, virus particles themselves are important sources of exogenous antigens feeding the immune system. It is nevertheless striking that other abundant virion proteins such as the large tegument protein BPLF1 and the major envelope protein BLLF1 (gp350)[32] were rarely if ever seen as targets for CD8+ memory responses in the present work or, for gp350, in earlier peptide screens[16, 44]. If the memory response is indeed re-shaped by cross-presentation, this implies that some virion proteins have differential access to the cross-priming pathway.

Importantly, the healthy carrier data move the overall picture of EBV-specific CD8+ T cell memory closer to that seen for HSV and CMV. Thus we detected responses to a mean of 14 different EBV lytic antigens per donor, compared to values of 8 and 17 in the HSV and CMV studies respectively[3, 4]. In total, healthy EBV carriers mounted responses against 30 of the 70 EBV lytic antigens tested, the corresponding values being 40/74 for HSV and 107/213 for CMV antigens. Of the 30 EBV lytic antigens identified, the two IE proteins remained frequent targets, accounting for almost 10% of all memory responses; this degree of focus, some three-fold above expectation based on IE sequence space, is very similar to that seen with the CMV IE proteins, though the trend is less marked with HSV. Apart from IE targets, detectable responses in the HSV and CMV studies were fairly evenly spread across early, early/late and true late antigens. However, particular antigens were again dominant; for example, among HSV early proteins the ribonucleotide reductase large subunit UL39 was a strong target[3], as was the large sub-unit BORF2 for EBV. Furthermore, among CMV and HSV late proteins some of the most frequently observed targets were abundant virion components [3, 4]; this is again in line with our own findings, although we noted interesting differences in detail. Thus the major capsid protein BcLF1 is the dominant L antigen for EBV-induced responses but its homologues in HSV (UL19) and CMV (UL86) are less frequently recognised. Conversely the most commonly recognised target in the CMV system is UL48, the large tegument protein, whereas its homologues in EBV (BPLF1) and HSV (UL36) seem to induce responses only rarely; for EBV the dominant tegument target is BNRF1, a protein unique to the gamma-herpesvirus subfamily which is delivered into cells by the virion and plays an important role in subverting cellular resistance to viral gene expression[45]. In the absence of corresponding data from other human herpesviruses, one does not know if these differences are sub-family-specific or unique to the individual agent.

Looking more broadly, the present paper brings our understanding of EBV lytic antigens as CD8+ T cell targets to the same level as that already established for latent antigens[8]. It shows that, while there are many lytic cycle proteins, they vary widely in immunogenicity and only a select few induce responses in a high proportion of people. Examples include BZLF1 (IE), BORF2 (E) and BcLF1 (L) that are seen by 11, 10 and 13, respectively, of our 14 study subjects. In addition the frequency of strong memory responses to another L antigen, BNRF1, is noteworthy because expression of this tegument component has also been detected in growth-transforming latent infection[16], one of several examples where detailed analysis of gene expression is beginning to blur the boundary between latent and lytic cycle[46, 47]. An improved understanding of lytic antigen targets is timely both because of this blurring and because recent evidence suggests that lytically-infected cells within foci of latent infection are critical for the efficient outgrowth of EBV-transformed B cells in vivo[48]. As a result, there is increasing interest in harnessing both latent and lytic antigen-specific T cells as effectors in two contexts: firstly in prophylactic vaccine design, where the aim is to contain the growth-transformed foci through which EBV colonises the general B cell system[36, 37], and secondly in targeting the EBV-transformed lymphoproliferative disease lesions to which T cell-compromised patients are especially prone[49]. It is hoped that the present study will aid progress towards those goals.

## MATERIALS AND METHODS

### Ethics Statement

The samples were from an already-existing study(Cellular immunity to herpesvirus infection: Studies with EBV & CMV). This study (14/WM/1254) was approved by West Midlands - Solihull Research Ethics Committee UK. All the samples were obtained following full written informed consent and were anonymized after collection.

### Donors

Patients in the acute phase on IM (within 7 days of the onset of symptoms) were identified on clinical grounds and diagnosis confirmed by high leukocyte counts and by IgM anti-virus capsid antigen (VCA) positivity. One patient also gave samples 4 years later, long after resolution of the illness. Other healthy carriers were individuals whom we knew from earlier serologic records had been infected with EBV for at least 5 years; none had a history of IM. Heparinised blood samples of 40-60ml were taken from both sets of donors. All donations were made in accordance with protocols approved by the South Birmingham Research Ethics Committee. For each donor, an aliquot of PBMCs was used to generate an EBV (B95.8 strain)-transformed LCL while the rest was used, either immediately or after cryostorage, as a starting population for the generation of effector preparations.

### HLA-expression constructs

DNA extracted from LCLs of donors was HLA-A,B,C-genotyped at the Anthony Nolan (www.anthonynolan.org) using sequence-specific oligonucleotide PCR. For generating HLA expression constructs, RNA was extracted from the LCL, reverse-transcribed (SuperScript III) and the specific HLA allotype amplified using specific primers. A-tails were added to synthesized HLA cDNA using Taq polymerase (30mins at 72 °C) and subsequently cloned into pCDNA3.1-V5/HIS TOPO expression constructs (ThermoFisher). Sequenced alleles were aligned against HLA sequences from the international ImMunoGeneTics project (www.imgt.org)

### EBV gene-expression constructs

EBV lytic genes were PCR-amplified from purified bacmid DNA derived from the B95.8 EBV strain[50] and were cloned into pCDNA3.1 vectors with the addition of a 5’ kozak sequence ‘GCCACC…’. In the case of two genes, BNLF2a and BNRF1, their coding sequences were separated into two fragments and individually cloned into expression constructs. Two lytic gene constructs, BBLF2/3 and BDRF1/BGRF1, were generated by fusing individual PCR products. Two other lytic gene constructs, for SM (BSLF2/BMLF1) and BPLF1, were kindly provided by Dr Shankar Swaminathan and Dr Maike Ressing respectively. A full list of the 70 lytic genes (as 72 constructs) is shown in Supplementary Figure 1.

### Generation of EBV lytic cell lysate

HEK293 cells stably transfected with B95.8 EBV strain as a bacmid [50] were maintained in standard culture medium (RPMI 1640 with 2mM L-glutamine (Sigma) containing 10% FCS, 500 IU/ml penicillin and 5000µg/ml streptomycin) supplemented with geneticin to maintain the EBV-bacmid. Lytic cycle entry was induced by transient transfection with the EBV-IE transactivator BZLF1. After 48 hours the cells were harvested, re-suspended to 4×10^6^ cells/ml in RPMI-1640 followed by 5 freeze-thaw cycles in a dry ice-ethanol bath before being sonicated for 30s. Samples were clarified by centrifugation at 1600rpm for 5mins before being stored at − 80°C.

### Primary effector preparations

Cryopreserved IM donor PBMCs were thawed and resuspended in standard culture supplemented with IL-2 as described previously[9]. The cells were then stimulated using gamma-irradiated and PHA-treated allogeneic PBMCs along with 30ng/ml anti-CD3 mAb (OKT3) and allowed to expand in culture for 2 weeks. The preparation was then depleted of CD4+ T cells (Dynabeads) and either used immediately or cryopreserved.

### Memory effector preparations

CD14+ monocytic cells were isolated from healthy carrier PBMCs using anti-CD14 coated beads (Miltenyi) and cultured for 4 days in standard culture medium containing 50ng/ml IL-4 and 50ng/ml GM-CSF (Peprotech)[51]. Remaining PBMCs were cryopreserved. The resultant immature monocyte-derived dendritic cells (moDCs) were harvested and re-suspended to 4×10^6^ cells/ml in standard culture media before exposure to the EBV lytic cell lysate at an equivalent cell ratio of 1:1. After 2 hours the mo-DC culture medium was supplemented with TLR-agonists RQR8-resiquimod (4µg/ml) and Polyinosinic-polycytidylic acid potassium salt (Poly(I:C)) (20µg/ml) as a maturation stimulus[52]. After 16-20 hrs the moDCs were harvested, washed and cultured with autologous lymphocytes cells at an effector: target ratio of 10:1. Specifically-activated cells were selected on the basis of CD137 upregulation as described in studies of HSV-specific T cell memory[3]. Note that we found CD137 upregulation to be optimal on day 3 of co-culture, longer than had been seen in the HSV system. This difference in timing probably reflects our use of cell-free extracts as an antigen source compared to the HSV study’s use of uv-irradiated lytically-infected cells[3]; dendritic cell cross-presentation is reportedly quicker for antigens acquired via phagocytosis of dead/dying cells[53–55]. Activated CD137+ CD8+ T cells were stained with Fixable Viability dye-eFluor 450 (ThermoFisher), anti-CD3-Allophycocyanin (APC), anti-CD8-APC-Cy7 and anti-CD137-PE (4-1BBL) (BD). Cells were sorted on a BD LSRFortessa X20, gating on live CD3+ lymphocytes and then CD8+CD137hi cells which were collected into tubes containing standard culture supplemented with IL-2. Sorted cells were centrifuged, re-suspended in 100µl standard culture supplemented with IL-2 and seeded into a 96 well U bottom plate. Using the same expansion conditions as with IM effectors, seeded cells were stimulated using irradiated and PHA-treated allogeneic PBMCs along with 30ng/ml anti-CD3 mAb (OKT3) and allowed to expand in culture for 2 weeks, then either used immediately or cryopreserved.

### Effector T cell screens

The target cells used in these were the EBV-negative cell lines MJS [17], for screening the full antigen panel, and COS7[3] for additional screening to restrict positive responses to a single HLA allele. Some 15,000 target cells were cultured per well in 96 well flat bottom culture plates. Duplicate wells were then transfected using Lipofectamine 2000 reagent (ThermoFisher) with 150ng/well of each individual EBV lytic gene construct plus 50ng of the donor’s paired HLA-A, −B or −C alleles. After 24 hours cells, transfected cells were trypsinized and transferred to 96 well V bottom plates. The cells were washed 2x within their wells through centrifugation and re-suspended in 100μl standard culture medium. Effector cell preparations from individual IM donors or healthy carriers were thawed and re-suspended in standard culture medium (without IL2) to a density of 1×10^6^ cells/ml. Those effectors were then seeded on top of transfected MJS targets at 1×10^5^ cells/ well in 100µl After a following 24 hours 50μl supernatant from each well was harvested and IFNγ production assayed in an IFNγ-capture ELISA. Significant responses were identified as those where IFNγ production was >1.7-fold above the background control value; this cut-off was selected on the basis of preliminary experiments as best discriminating between weak but reproducible positive responses and negative responses. Where significant recognition was observed, the effectors were re-tested against the same EBV lytic gene construct expressed in COS7 cells alongside the relevant HLA-A, B or C alleles now tested as individual restricting elements. Note that, while COS7 is of simian origin, MJS is a human melanoma-derived cell line and expresses HLA-A*0101, B*0801 and C*0701 endogenously; in the screening assays, however, antigen presentation through MJS’s endogenous alleles was minimal and positive recognition required the presence of the exogenous HLA constructs.

### Statistical analyses

For univariate analyses, where data did not follow a normal distribution Mann-Whitney U test (for unpaired data) was used. Where data followed a normal distribution, it was analysed by two-tailed t-test. Where there were multiple independent variables, they were categorised and then analysed by Kruskal-Wallis test with a Dunn’s multiple comparison. The correlation of immunogenicity and antigen size was carried out using Linear Regression analysis. All analyses were performed with Prism version 6.0 (Graphpad software, San Diego, USA).

## Acknowledgements

We thank the Dr Claire Shannon-Lowe for assistance with culture and induction of HEK293 cells stably transfected with B95.8 EBV strain as a bacmid and Drs Shankar Swaminathan and Maike Ressing for the SM (BSLF2/BMLF1) and BPLF1 gene constructs.

## Supporting Information Legends

**Supplementary Figure 1:** Individual results from proteome-wide screening of the primary CD8+ T cell response in IM patients IM217, IM223, IM239, IM243, IM 249 and IM269. Data are presented as in Figure 2.

**Supplementary Figure 2:** Individual results from proteome-wide screening of the primary CD8+ T cell response in healthy carriers HC2, HC3, HC4, HC5, HC6 and HC7. Data are presented as in Figure 2.

**Supplementary Table 1:** EBV lytic cycle genes cloned into expression vectors. Footnote: The genes are arranged in IE, E and L categories and shown alongside the function (where known) and size (number of amino acid residues) of their protein products. * BHLF1 and LF3 contain many repeat domains and their sizes have been reduced to reflect their unique amino acid content. ** Two genes, BNLF2a and BNRF1, were expressed as separate N-terminal and C-terminal fragments.

**Supplementary Table 2:** Immunogenic EBV antigen/HLA allele combinations identified in this work Footnote: Data are compiled for all the individual HLA-A, HLA-B and HLA-C alleles possessed by study donors. For each allele, the Table records the number of allele-positive donors, those antigens through which an allele-restricted response was observed, and the donors who made such a response. Shading identifies “prevalent” responses, i.e cases where more than half the donors with the relevant allele made a response to the specific allele/antigen combination.

